# Accurate determination of house dust mite sensitization in asthma and allergic rhinitis through cytometric detection of Der p 1 and Der p 2 binding on Basophils (CytoBas)

**DOI:** 10.1101/2023.08.15.553357

**Authors:** Lin Hsin, Nirupama Varese, Pei Mun Aui, Bruce D. Wines, Laurent Mascarell, Mark P. Hogarth, Mark Hew, Robyn E. O’Hehir, Menno C. van Zelm

**Author notes:** **Corresponding Author:** Menno C. van Zelm, PhD, Allergy and Clinical Immunology Laboratory, Monash University, 89 Commercial Rd, Melbourne, VIC 3004, Australia. **Email:**.

## Abstract

**Background:** House dust mite (HDM) is the commonest allergen trigger globally for allergic rhinitis and atopic asthma. To expedite accurate confirmation of allergen sensitization, we designed fluorescent allergen tetramers to directly stain specific IgE on basophils to detect allergen sensitization using the flow cytometric CytoBas assay.

**Methods:** Recombinant proteins of major HDM allergens (component), Der f 1, Der p 1 and Der p 2 were biotinylated and conjugated with fluorochrome streptavidins as tetramers. Blood samples from 64 HDM-allergic patients and 26 non-HDM-sensitized controls were incubated with allergen tetramers for evaluation of basophil binding (CytoBas) and activation (BAT) with flow cytometry.

**Results:** The tetramers effectively bound and activated basophils from allergic patients but not non-sensitized controls. CytoBas with Der p 1 as a single allergen had comparable sensitivity and specificity (92% and 100%) to BAT (91% and 100%), similarly for CytoBas with a single Der p 2 (95% and 96%) to BAT (95% and 87%) in detecting allergen sensitization. A positive staining for Der p 1 and/or Der p 2 was 100% sensitive and 96% specific for HDM allergy.

**Conclusions:** CytoBas has diagnostic accuracy for group 1 and group 2 HDM allergens that is comparable to a BAT assay, but with additional advantages of multiple allergen components in a single tube and no requirement for *in vitro* basophil activation. These findings endorse a single, multiplex CytoBas assay for accurate and component-resolved diagnosis of aeroallergen sensitization in patients with allergic asthma and/or rhinitis.

**CAPSULE SUMMARY:** A single flow cytometry stain of basophils (CytoBas) with both Der p 1 and Der p 2 provides >95% specificity and sensitivity for detection of functional HDM allergen sensitization.

**Highlights:**

- Fluorescent tetramers of recombinant Der f 1, Der p 1 and Der p 2 can be used to detect functional IgE sensitization to house dust mite (HDM) by flowcytometric detection on basophils (CytoBas).
- A single CytoBas assay with inclusion of both Der p 1 and Der p 2 can detect HDM sensitization with >95% sensitivity and specificity.

## INTRODUCTION

Atopic asthma and allergic rhinitis (AR) affect about 300 million people worldwide. Impacting about 50% of these individuals, house dust mite (HDM) is the most common allergic trigger globally.^1^ Disease control typically relies on allergen avoidance where possible, which is challenging, given the ubiquity of HDM.^2^ Pharmacotherapy is effective but requires regular administration.^3^ Allergen immunotherapy (AIT) is the only disease-modifying treatment for HDM allergy, with the potential for a sustained response beyond the treatment period.^4^ There is thus a compelling need for early and accurate confirmation of suspected HDM allergen sensitization^5^ to facilitate optimal treatment.^6^

Conventional diagnostics for determining atopic sensitization in allergic asthma and rhinitis include skin prick testing (SPT) and allergen-specific serum IgE detection.^7^ However, their drawbacks include poor specificity (71-76% for SPT; 52-58% for serum IgE detection), partly due to the frequent use of allergen extracts that contain non-allergenic components.^8–10^

Additional diagnostic modalities are available. The basophil activation test (BAT) is a functional assay used in some research laboratories, through which the specificity of effector cell-bound IgE is evaluated.^11^ *In vitro* exposure to allergens induces cross-linking of the high-affinity FcεRI on specific IgE-loaded blood basophils.^12^ Basophil activation leads to degranulation and surface expression of CD63, which is typically used as an activation marker in BAT.^13, 14^ Frequencies of CD63 positive basophils correlate better (r=0.5) with clinical severity than SPT or sIgE levels (r=0.24 and r=0.33, respectively).^15, 16^ However, BAT is labor intensive– requiring fresh blood, careful allergen titrations, and time-frame restrictions – which hinders its implementation into routine clinical practice.^17^

Most recently, component-resolved diagnostics (CRD) are being applied to determine allergen sensitization by detecting IgE specific to allergen components, which can be either purified from natural allergen sources or recombinantly produced.^18^ To date, 76 HDM allergen components across various species have been defined by the WHO/IUIS Allergen Nomenclature Sub-committee.^19^ IgE responses to some of these can be cross-reactive,^20^ including tropomyosin, which is highly homologous among invertebrates, such as insects (cockroaches), mollusks and crustaceans (e.g. shrimp).^21^

There are 2 common HDM species globally, which coexist in many geographical regions, although *Dermatophagoides pteronyssinus* (Der p) is more commonly found in Europe and Australia, and *D. farinae* (Der f) in the USA and Asia.^22, 23^ Fecal pellets are the main source of allergens,^24^ with group 1 and group 2 allergens being the most immunodominant proteins; 95% of HDM-allergic patients have specific IgE against either or both.^25^ The group 1 allergens, Der p 1 and Der f 1, are cysteine proteases, which have >90% protein sequence homology, resulting in high IgE cross-reactivity among allergic patients.^26^ Der f 2 and Der p 2 are lipid-binding proteins with 87% sequence homology.^27, 28^

CRD is gaining utility in both research and clinical settings and can be highly beneficial for HDM allergy management. ^29^ Detection of specific IgE to Der p 1 and Der p 2 can identify 75-85% of allergic patients,^30, 31^ with a better positive predictive value for HDM allergy than HDM extract. CRD can distinguish clinically-relevant allergen sensitization to HDM components from cross-reactivity with shrimp allergens^32, 33^ allowing optimal treatment prescription for HDM allergic patients.^34^

We recently developed a CRD which accurately detected functional sensitization to bee venom (Api m 1) and RGP (Lol p 1 and Lol p 5), using recombinant allergen components for multiplex cytometric staining of blood basophils (CytoBas) without the need for their activation as required by BAT.^35, 36^ In this study, we explored the diagnostic performance of CytoBas using Der f 1, Der p 1 and Der p 2 to detect sensitization to HDM, the commonest trigger for perennial atopic asthma and allergic rhinitis globally.

## METHODS

### Study Participants and sample collection

HDM allergic participants (n=64) were recruited from the Allergy Clinic at the Alfred Hospital in Melbourne, Victoria, Australia on the basis of a diagnosis of perennial allergic rhinitis with or without asthma and serum HDM-specific IgE of ≥0.35kU_A_/L (ImmunoCAP, Phadia, Uppsala, Sweden). Non-sensitized controls (n=26) were recruited based on the absence of HDM sensitization, as evidenced by the absence of clinical symptoms and a negative skin prick test (<1mm) or serum HDM-specific IgE levels of <0.35 kU_A_/L (ImmunoCAP). Exclusion criteria for all subjects were immunodeficiency, gastrointestinal complications, AIT within the last five years, or prescribed beta-blockers or oral corticosteroids. The use of symptom relievers for allergic rhinitis, such as antihistamines and topical corticosteroids, was allowed. Lithium-heparinized blood samples were collected from all subjects and processed within 24 hours for flow cytometry (see below) serum and PBMC isolation and cryopreservation. The study was conducted in compliance with the Declaration of Helsinki, and written informed consent was obtained from each participant prior to inclusion (Alfred Ethics Project ID: 297-20).

### Recombinant protein production

Protein sequences of Der f 1.0101, Der p 1.0102, and Der p 2.0101 were obtained from the Allergen Nomenclature website (allergen.org).^37, 38^ Full length Der f 1.0101 and Der p 1.0102 constructs were generated with the native leader sequence for secretion, a BirA tag for biotinylation, and a 6-His tag for cobalt column binding during protein purification (**Figure 1**). Both constructs contained the C35S mutation to disrupt their cysteine protease activity.^39^,

**Figure 1.**
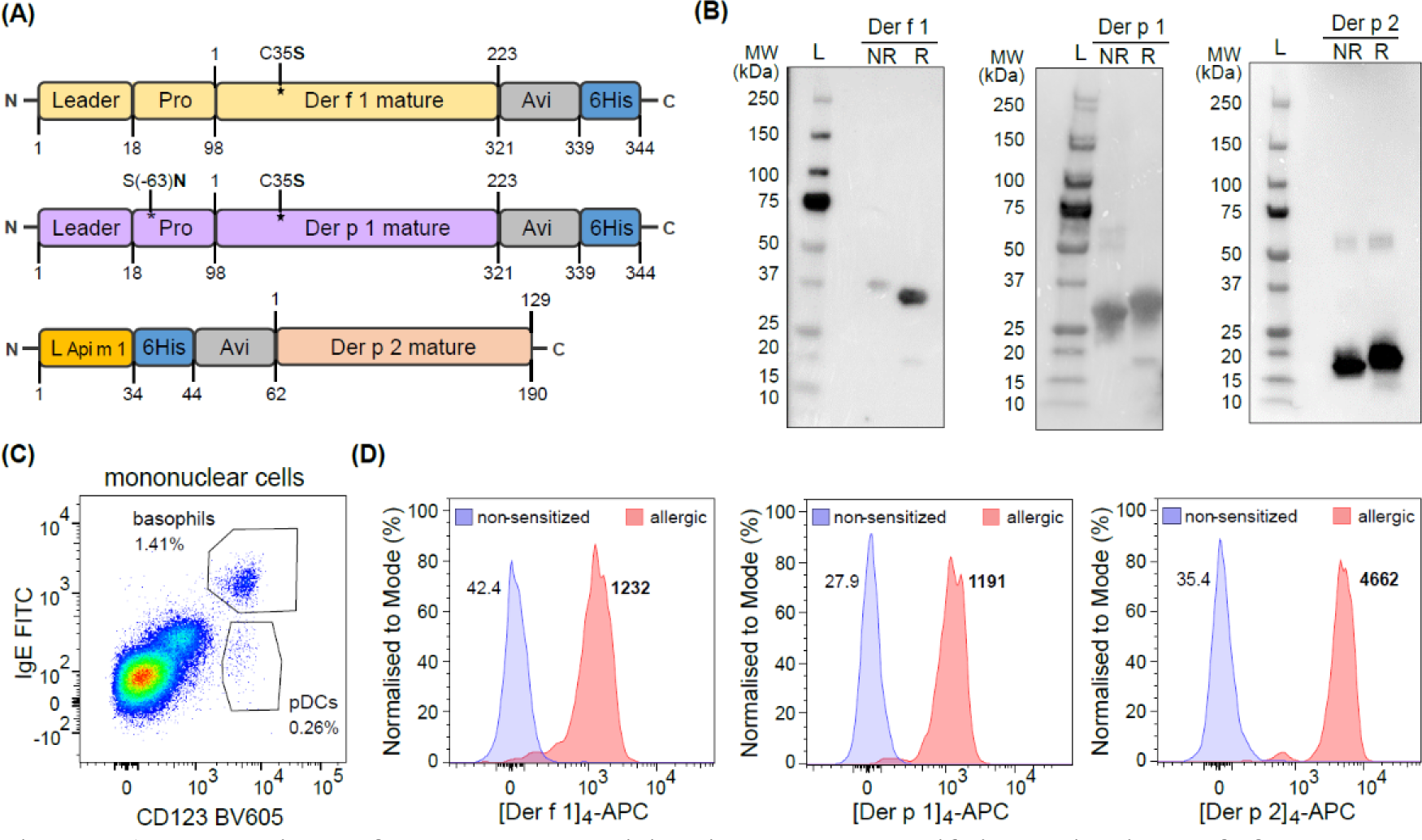
Detection of allergen sensitization by quantifying binding of fluorescent allergen components of HDM to basophils. **(A)** Schematic diagram of DNA constructs for Der p 1 and Der p 2 protein generation. **(B)** Anti-His western blots of recombinant allergens in non-reduced (NR) and reduced (R) forms. Protein ladders (L) are shown to ensure proteins possess the molecular weight (MW) of the sequence-predicted sizes, Der f 1 (37kDa); Der p 1 (37kDa); and Der p 2 (21kDa). **(C)** Within gating of circulating basophils from mononuclear cells, **(D)** representative flow cytometry plot of tetramer staining by [Der f 1]_4_-APC, [Der p 1]_4_-APC and [Der p 2]_4_-APC, respectively, between non-sensitized and allergic individuals.

^40^ Additionally, the Der p 1 construct contained an S(−63)N mutation in the prodomain to disrupt the N-glycosylation motif (NKS from codons −65 to −63). This motif is absent from Der f 1, and its presence in Der p 1 was previously shown to impair efficient protein production.^41^ The Der p 2.0101 construct was generated with an N-terminal leader sequence from Api m 1 (*Apis mellifera*),^35^ a 6-His tag and a BirA sequence (**Figure 1**). All three constructs were codon-optimized for *Spodopera frugiperda* (Sf) and cloned into the pFastBac vector (Thermo Fisher Scientific, Waltham, MA, USA), before incorporating into a Bacmid for baculovirus production.^35^ Bacmids with the construct of interest were transfected into Sf21 cells, which were then cultured in a shaker incubator at 27°C. Following viral amplification, the supernatant was harvested following centrifugation. The 6-His tagged recombinant proteins were purified from the supernatant by gravity-fed through TALON® Metal Affinity Resin (Takara Bio, Shiga, Japan). Following a wash step with PBS (pH 8), the proteins were eluted with imidazole (200mM in PBS, pH 8). The eluate was concentrated with an ultra-0.5 centrifugal filter unit (Amicon^®^, Merck KGaA, Darmstadt, Germany) at 3000 *g for 30 mins at 4°C, followed by dialysis and concentration in 10mM TRIS (pH 8). The purified recombinant proteins were subjected to targeted-biotinylation by overnight incubation at room temperature (RT) with a BirA enzyme (2.5µg/mL) in 10mM TRIS containing 62.5 mM Bicine-HCl, pH 8.3; 62.5 µM D-Biotin; 12.5 mM ATP; and 12.5 mM MgOAc, followed by dialysis against 10mM TRIS (pH 8). The biotinylated proteins were tetramerized by adding APC-conjugated streptavidin (Thermo Fisher Scientific) at an allergen: streptavidin molar ratio of 4:1.^35, 42^

### Western blotting

Purified recombinant His-tagged proteins were diluted to 0.5µg/ml in either Laemmli sample buffer alone (Bio-Rad, Hercules, CA, USA) or containing 50 mM dithiothreitol (DTT) (reducing buffer) and treated at 90°C for 10 min. The samples and a protein ladder (a 1:1 mix of All Blue Prestained Protein Standards: Unstained Protein Standards; Precision Plus Protein™, Bio-Rad) were loaded to 4-15% mini-PROTEAN™ TGX Stain-Free™ Protein Gels (Bio-Rad), and separated at 200V for 30 mins. The protein gel was exposed and imaged on ChemiDoc Imager (Bio-Rad) prior to protein transfer onto a PVDF membrane (Bio-Rad) using the Trans-Blot Turbo Transfer System (Bio-Rad). The membrane was blocked with 3% skin-milk-powder in Tris-buffered saline (Bio-Rad) with 0.1% Tween^®^ 20. Protein detection was performed by incubation with a 6x-His Tag mouse monoclonal antibody (clone: HIS.H8; Thermo Fisher Scientific), followed by a secondary goat-anti-mouse HRP antibody (clone NA9310 N; Cell signaling Technology, Danvers, MA, USA) and incubation with the Luminata forte western HRP substrate (Merck Millipore, Burlington, MA, USA). Colorimetric (standards) and chemiluminescent detection (His-tagged proteins) were performed using the ChemiDoc Imager (Bio-Rad).

### Basophil activation test (BAT) and basophil staining for CytoBas

BAT and CytoBas were performed on fresh whole blood samples collected from all study subjects within 24hr of collection, as described previously.^35^ The circulating basophils were first primed with IL-3 by incubating for 10mins at 37°C with stimulation buffer (HEPES 20 mM, NaCl 133 mM, KCl 5 mM, CaCl_2_ 7mM, MgCl_2_ 3.5 mM, BSA 1 mg/mL, rIL-3 2 ng/mL and Heparin 20 µL/mL, pH 7.4). Cells were then stimulated with HDM extract (0.001, 0.01, 0.1, 1 µg/mL; Stallergenes Greer, London, UK), stimulated, and directly stained with fluorescent allergen tetramers (0.01, 0.1, 1 µg/mL), or streptavidin conjugate only (1 µg/mL; Allophycocyanin (APC) conjugate, Thermo Fisher Scientific) for 20 mins at 37°C. Alongside, stimulation with rabbit anti-human IgE (Dako, Carpinteria, CA, USA) and fMLP (N-Formylmethionine-leucyl-phenylalanine; Sigma Aldrich, St. Louis, MO, USA), respectively, were used as positive controls for FcεRI-dependent and FcεRI-independent basophil degranulation. Activation was halted by incubating samples on ice for 5 mins. The cells were then washed with cold wash buffer (HEPES 20mM, NaCl 133mM, KCl 5mM and EDTA 0.27 mM, pH 7.3) before staining surface markers for flow cytometry.

### Flow cytometry

Following basophil activation or direct staining, the basophils were incubated with IgE-FITC (polyclonal, Thermo Fisher Scientific) and/or HLA-DR-APC (L243; BioLegend, San Diego, CA, USA), as well as CD123-BV605 (6H6; BioLegend), Fixable live/dead dye 700 (BD Biosciences, San Jose, CA, USA) and CD63-PE (H5C6, BD Biosciences), on ice for 20 mins. Then, the samples were incubated in red-blood-cell lysis buffer (NH_4_Cl 155 mM, KHCO_3_ 9.9 mM and EDTA 1mM) for 10 min at RT and washed with wash buffer (HEPES 20mM, NaCl 133mM, KCl 5mM and EDTA 0.27 mM, pH 7.3). After that, samples were incubated in paraformaldehyde (PFA; 1%; ProScitech, QLD, Australia) for 20 mins at RT and then fixed samples were then resuspended in 100µL wash buffer prior to acquisition on the 4-laser BD^®^ LSRII (BD Biosciences). Instrument setup and calibration were performed using EuroFlow standard operation procedures as described previously.^43^

### Data analysis and statistics

Flow cytometry data were analyzed with FACS DIVA V6.3.1 software (BD Biosciences) and FlowJo v10.8.0 software packages (FlowJo LLC, Ashland, OR, USA). Statistical analyses were performed using GraphPad Prism (v9.0.1). The non-parametric Wilcoxon signed-rank test was used for paired data, and the non-parametric Mann-Whitney U-test was for non-paired data. Receiver operating characteristic (ROC) curves were utilized to determine the performance capabilities (specificity and sensitivity) of allergen components in the BAT and CytoBas assays between HDM allergic patients and HDM-non-sensitized. P-values of <0.05 were considered statistically significant.

## RESULTS

### Design and production of recombinant allergen tetramers for CytoBas

Production of full-length recombinant Der f 1 with the C35S mutation and C-terminal 6-His and BirA tags was successful (**Figure 1A**). Furthermore, introduction of the S(−63)N mutation in Der p 1, to disrupt the N-glycan attachment at N-65, and model the lack of a glycosylation motif in Der f 1, resulted in successful production of this recombinant protein. The full-length Der p 2 protein was generated with the N-terminal Api m 1 leader sequence,^35^ 6-His, and BirA tags (**Figure 1A**). Purified recombinant allergens were detected at the expected molecular weights by western blotting using an anti-6-His antibody (ThermoFisher) (**Figure 1B**). Following targeted biotinylation, these proteins were tetramerized with streptavidin-APC for BAT and CytoBas assays.

To confirm the ability of each recombinant allergen protein tetramer to bind to IgE on basophils by flow cytometry, we first gated on IgE^+^CD123^+^ basophils (**Figure 1C**). The APC-conjugated [Der f 1]_4_, [Der p 1]_4_, and [Der p 2]_4_ positively stained basophils derived from HDM-allergic patients but not those from non-sensitized controls (**Figure 1D**).

### Recombinant Der f 1, Der p 1, and Der p 2 tetramer staining can detect functional HDM-allergen sensitization

The ability of [Der f 1]_4,_ [Der p 1]_4_, and [Der p 2]_4_ to detect functional allergen sensitization was determined by measuring the allergen binding (**Figure 2A, B**) and activation (CD63 positivity; **Figure 2C, D**) using blood samples from 62 HDM-allergic patients and 26 non-sensitized controls (**Table 1**). Detection of blood basophils post-activation was performed with the gating strategy shown (**Suppl. Figure 1**), and plasmacytoid dendritic cells (pDC) were gated as a known negative control.^35^ Within the gated basophils, the median fluorescent intensity (MFI) of the APC-allergen tetramers was examined as the allergen-specific IgE binding intensity, and CD63 expression was used to investigate activation and degranulation. As controls for basophil activation, separate samples from each donor were incubated with serial dilutions of HDM extract, anti-IgE or fMLP, a bacterial tripeptide that activates basophils through a G-protein coupled receptor, independent from Fc□RI.^17^

**Figure 2.**
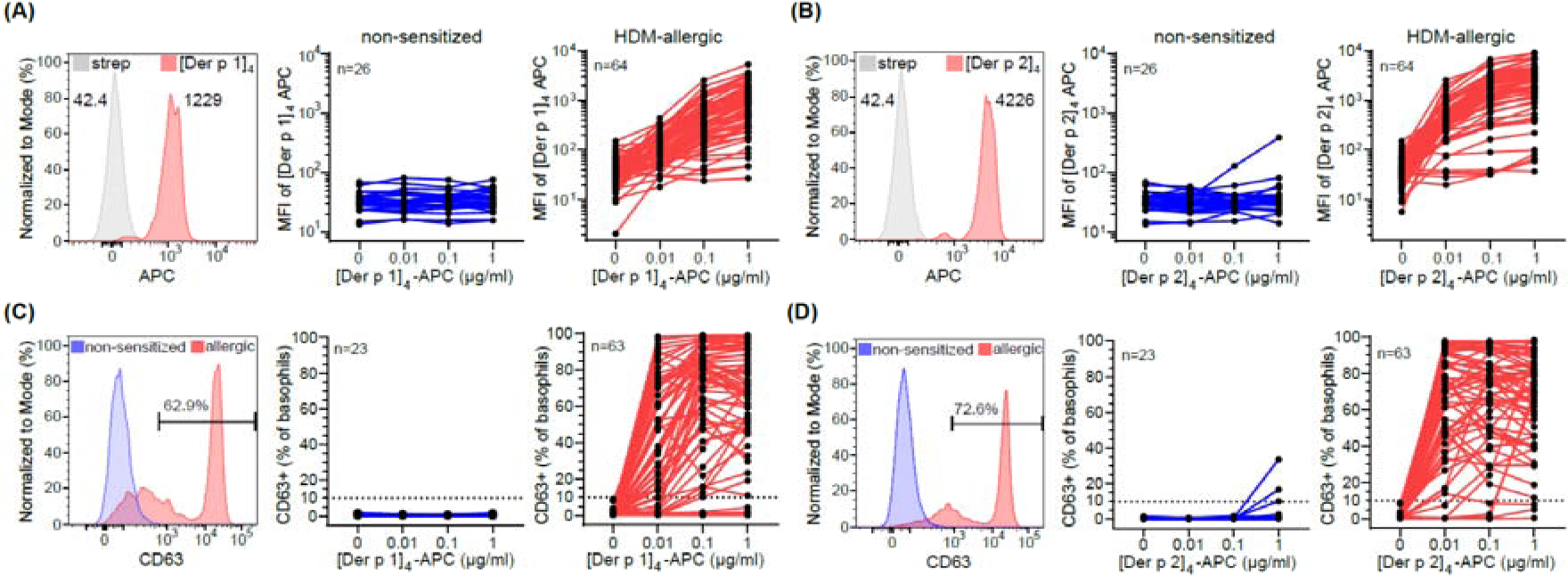
Recombinant Der p 1 and Der p 2 bind to basophils from HDM-allergic patients in a concentration-dependent manner and induce basophil activation. **(A)/(B)** Representative staining of basophils from an HDM-allergic patient blood sample, incubated with [Der p 1]_4_-APC/ [Der p 2]_4_-APC or streptavidin-APC. Median fluorescence intensity (MFI) of basophils from non-sensitized controls and HDM-allergic subjects after incubation with 1μg/mL streptavidin-APC, presented as the baseline control “0 μg/mL” or 0.01, 0.1 and 1μg/mL [Der p 1]_4_-APC/ [Der p 2]_4_-APC. **(C)/(D)** Representative histograms of CD63 expression on basophils and frequencies of basophils from non-sensitized and HDM-allergic subjects following stimulation with streptavidin-APC or ascending concentrations of [Der p 1]_4_-APC/ [Der p 2]_4_-APC.

**Table 1.**
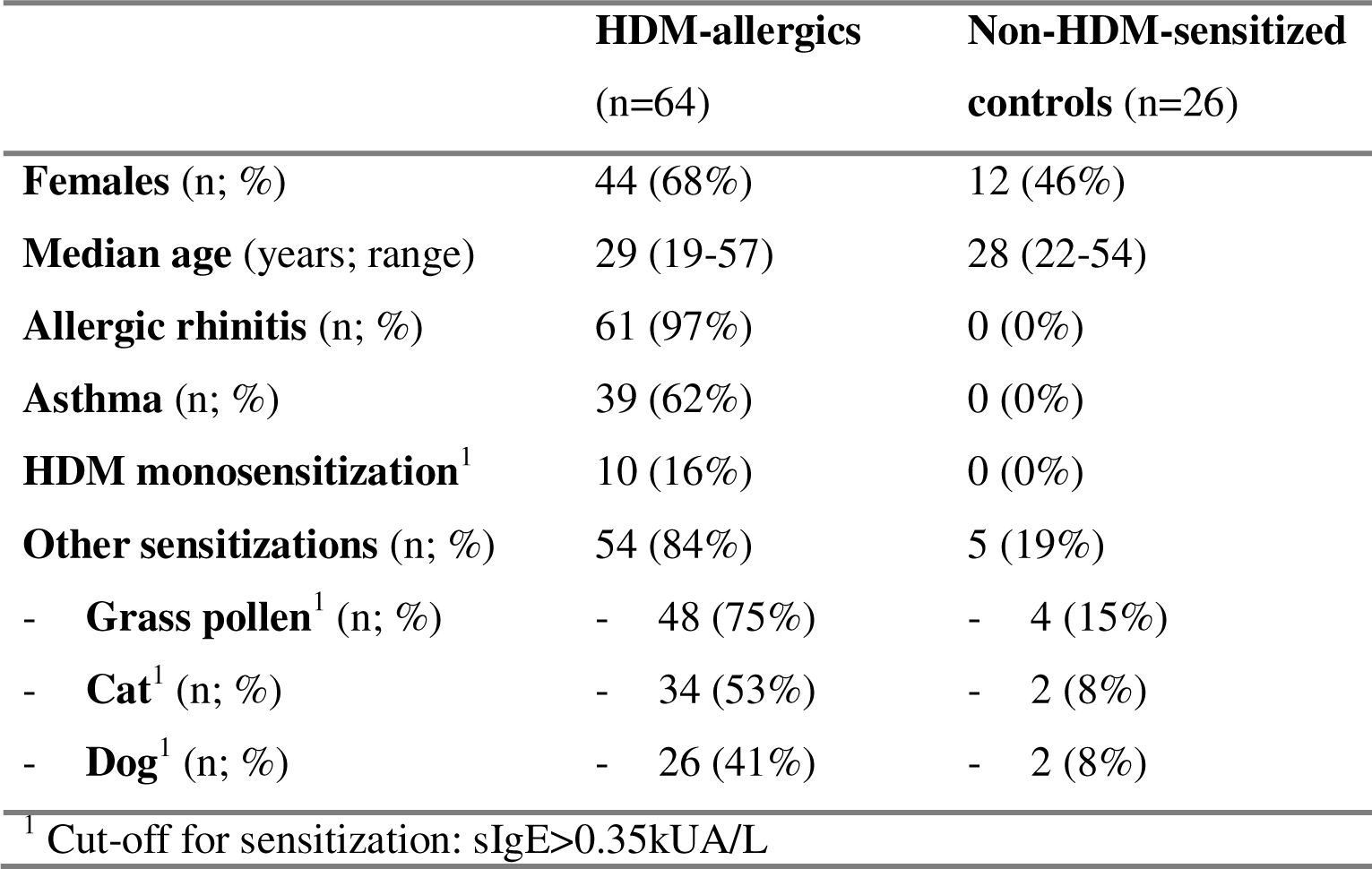
Demographics and characteristics of study participants.

Following HDM extract incubation, basophils from HDM-allergic patients only showed an increased expression of CD63^+^ (**Suppl. Figure 2A**). Basophils from both HDM-allergic (n=62/64) and non-HDM-sensitized (n=25/26) individuals expressed CD63 after stimulation with the IgE-independent stimulus fMLP (**Suppl. Figure 2B**). Anti-IgE stimulation induced CD63 expression on the basophils of most HDM-allergics (n=60/64) and control subjects (n=23/26) (**Suppl. Figure 2C**). Donors whose basophils did not respond to both anti-IgE and HDM (extract or components) were classified as non-responders and excluded from further BAT analysis (n=1 allergic patient; n=3 controls).

While there was no detectable tetramer staining in nearly all the controls, a concentration-dependent increase in MFI was evident in the HDM-allergic group following incubation with either [Der f 1]_4_ (**Suppl. Figure 3**), [Der p 1]_4_ (**Figure 2A**) or [Der p 2]_4_ (**Figure 2B**). The increased staining intensity was accompanied by an increase in CD63^+^ basophil frequencies in allergic patients but not controls (**Figure 2C, D**).^17^ Basophils from all subjects were positive for IgE, whereas pDC were not (**Suppl. Figure 4**). In line with all basophils being IgE^+^ in sensitized patients, all basophils showed positive staining for the allergen tetramers. CD63^-^ and CD63^+^ basophils from all HDM component-specific allergic patients showed significant (p<0.0001) binding with either tetramer, compared with streptavidin-APC control. More importantly, CD63^-^ and CD63^+^ basophils of allergic individuals were found to bind [Der p 2]_4_ to a similar extent (**Suppl. Figure 5**). Thus, despite not all basophils from a single individual undergo *in vitro* degranulation upon allergen binding, basophils have a similar capacity to bind allergen with specific IgE. These findings confirmed our tetramers were being recognized and can detect HDM sensitization in allergic patients.

### High diagnostic capacity to detect HDM allergy sensitization with basophil staining

To evaluate the diagnostic ability of CytoBas in comparison to BAT for diagnosing HDM allergen sensitization, we compared results between the non-sensitized controls and allergic patients, and ROC curves generated from both groups were analyzed.

Tetramer staining intensities were determined by calculating the ratio between MFI of allergen tetramer (1µg/mL) and streptavidin-APC staining. The [Der p 1]_4_ staining ratio was >2 for 59/64 allergic patients, compared to <2 for all 26 control subjects (**Figure 3A**). Basophil activation with 1µg/mL [Der p 1]_4_ induced >10% CD63^+^ basophils in 57/63 patients, whereas this population was <10% in all evaluable controls (n=23) (**Figure 3B**). In a direct comparison, the [Der p 1]_4_ CytoBas had comparable sensitivity (92%) to BAT (91%) for detecting HDM sensitization, while both tests demonstrated 100% specificity (**Table 2**). In addition, ROC curve analyses showed very high AUC for both [Der p 1]_4_ CytoBas (0.9549) and [Der p 1]_4_ BAT (0.9683).

**Figure 3.**
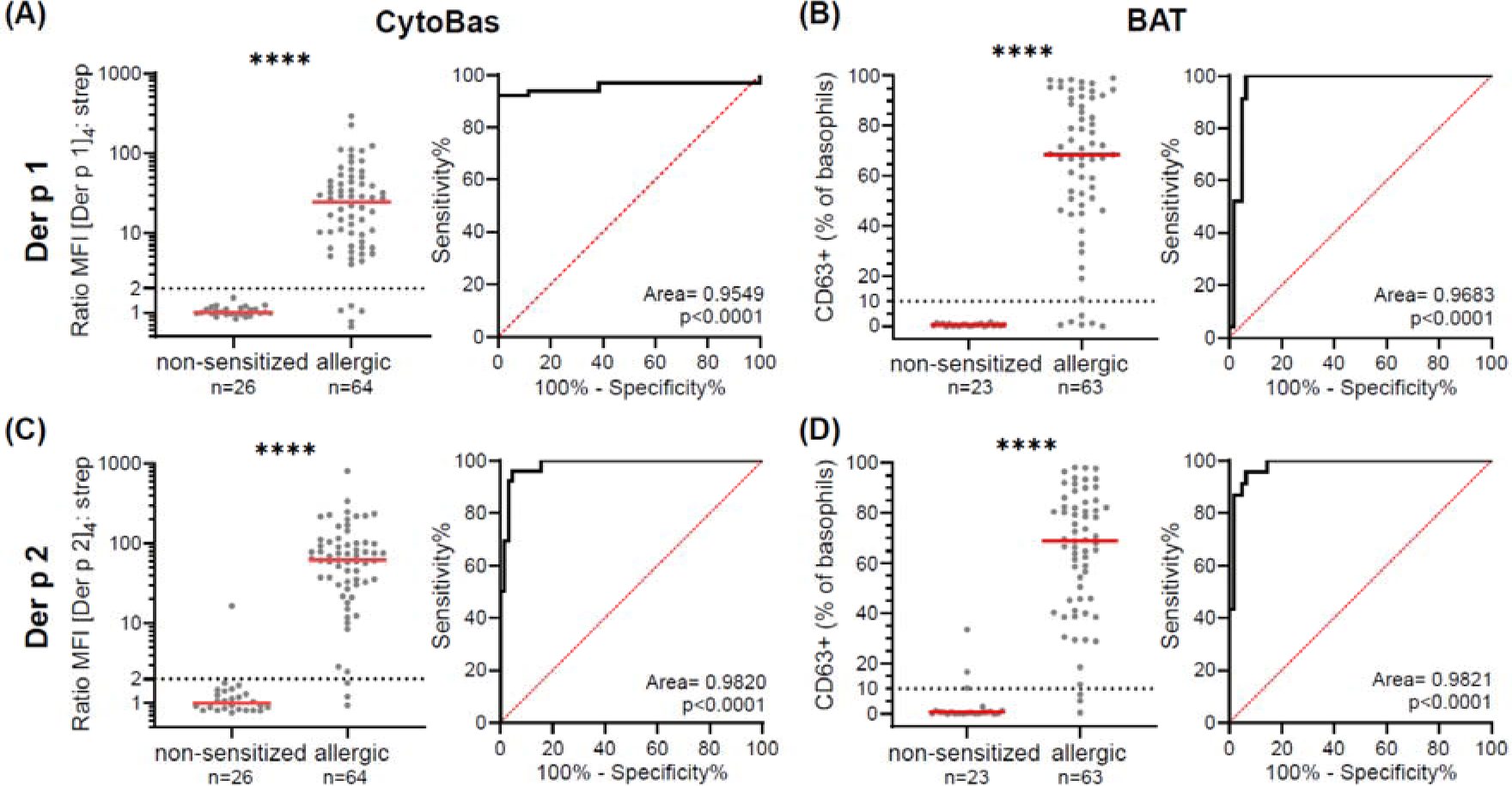
The sensitivity and specificity of CytoBas are comparable to BAT for HDM allergy detection. Staining intensities of HDM tetramers in CytoBas via the ratio of **(A)** [Der p 1]_4_-APC at 1μg/mL/streptavidin-APC, **(C)** [Der p 2]_4_-APC at 1μg/mL/streptavidin-APC and respective ROC on staining intensity. Frequency of CD63^+^ basophils in BAT following *in vitro* stimulation with 1µg/mL of **(B)** [Der p 1]_4_-APC, **(D)** [Der p 2]_4_-APC and receiver operator characteristics (ROC) of CD63 positivity. Statistics: Mann-Whitney U test; for ROC, Wilson/Brown method for confidence level (>95%) determination. **** p <0.0001.

**Table 2.** Diagnostic performance of basophil activation test (BAT) and CytoBas assays with HDM components.

| Assay | HDM Component | True Positive (n) | False Positive (n) | False Negative (n) | True Negative (n) | Sensitivity (%) | Specificity (%) | Accuracy (%) |
| --- | --- | --- | --- | --- | --- | --- | --- | --- |
| BAT | Der f 1<br>(35 patients, 15 controls) | 27 | 0 | 8 | 15 | 75.0 | 100 | 82.4 |
|  | Der p 1<br>(63 patients, 23 controls) | 57 | 0 | 6 | 23 | 90.5 | 100 | 93.0 |
|  | Der p 2<br>(63 patients, 23 controls) | 60 | 3 | 3 | 20 | 95.2 | 87.0 | 93.0 |
|  | **Der p 1 and/or Der p 2**<br>(63 patients, 23 controls) | **62** | **3** | **1** | **20** | **96.9** | **86.9** | **94.2** |
| CytoBas | Der f 1<br>(36 patients, 17 controls) | 32 | 0 | 4 | 17 | 88.9 | 100 | 92.5 |
|  | Der p 1<br>(64 patients, 26 controls) | 59 | 0 | 5 | 26 | 92.2 | 100 | 94.4 |
|  | Der p 2<br>(64 patients, 26 controls) | 61 | 1 | 3 | 25 | 95.3 | 96.2 | 95.6 |
|  | **Der p 1 and/or Der p 2**<br>(64 patients, 26 controls) | **64** | **1** | **0** | **25** | **100** | **96.2** | **98.9** |

The [Der p 2]_4_ staining ratio was >2 in 61/64 allergic patients, and <2 in 25/26 control subjects (**Figure 3C**). Basophil activation with 1µg/mL [Der p 2]_4_ induced >10% CD63^+^ basophils in 60/64 patients, and this population was <10% in 20/23 controls (**Figure 3D**). The sensitivity and specificity (95% and 87%) of [Der p 2]_4_ CytoBas were similar to BAT’s (95% and 96%) (**Table 2**). The AUC from the ROC curve analyses demonstrated enhanced performance compared to [Der p 1]_4_: 0.9821 for [Der p 2]_4_ CytoBas and 0.9820 for [Der p 2]_4_ BAT.

A limited set of samples were run with Der f 1 (**Suppl. Figure 3**), as this is not the major sensitizer in Australia.^22^ Nevertheless, the [Der f 1]_4_ staining ratio was >2 for 32/36 allergic patients, and <2 for all 17 control subjects (**Suppl. Figure 3A-E**). Only 27/35 patients showed >10% CD63^+^ basophils following stimulation with 1µg/mL [Der f 1]_4_, and all the 15 evaluable controls had <10% stimulation (**Suppl. Figure 3F-J**). CytoBas with [Der f 1]_4_ yielded a very similar sensitivity and specificity as [Der p 1]_4_ CytoBas, and large AUC in the ROC analysis (0.9379) (**Suppl. Figure 3E**), underlining the high cross-reactivity due to the great sequence homology between the two proteins.^44^

### Multiplex component-resolved diagnosis with the inclusion of both Der p 1 and Der p 2

Our CytoBas approach enables the inclusion of multiple allergen components in a single assay. ^35^ To evaluate the potential of combined evaluation with two allergen components, we depicted the allergen: streptavidin staining ratios of [Der p 1]_4_ versus [Der p 2]_4_ (**Figure 4A**). Detection of positive sensitization to HDM was defined as one or both MFI ratios above 2. Thus, only datapoints within the lower left quadrant were defined as negative. All 64 HDM-allergic patients were positive for one or both allergens, and 25/26 control subjects were below the cut-off ratio of 2 for both allergens. Despite basophil activation requiring separate tubes and multiple dilution procedures to evaluate each component, a direct comparison was made with BAT (**Figure 4B**). This showed that 1/63 allergic patients was negative for both components, and 3/23 control individuals were positive for Der p 2 (**Table 2**). Thus, by using two allergen components, the CytoBas approach achieved 100% sensitivity and (96%) specificity. The BAT was equally sensitive (100%), but less specific (87%). This analysis demonstrated the capacity for CytoBas to enable CRD in a single tube for differential detection of monosensitization to either Der p 1 (3/64) or Der p 2 (5/64), or both (56/64) in HDM-allergic patients.

**Figure 4.**
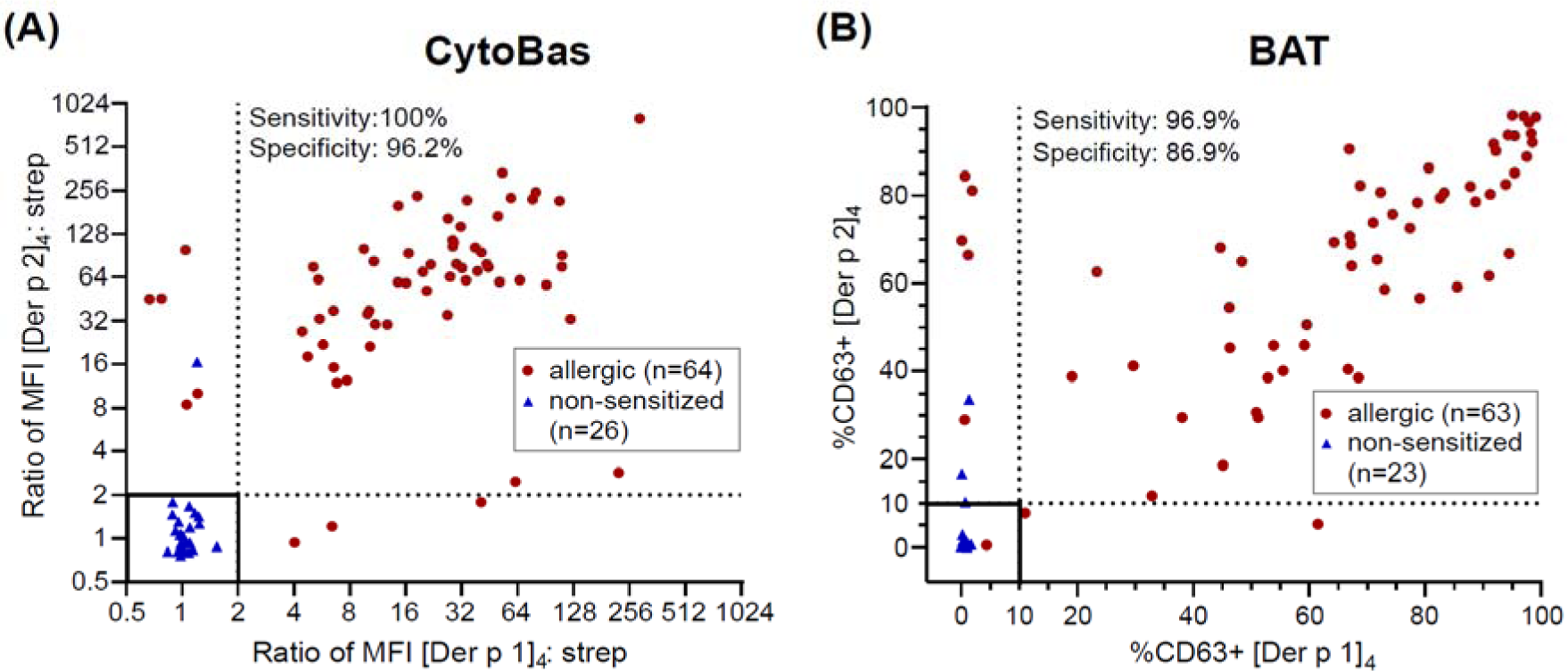
Combined evaluation of Der p 1 and Der p 2 tetramers for component-resolved diagnostic tool for HDM allergy. **(A)** The capacity of CytoBas to detect HDM sensitization was evaluated based on positive [Der p 1]4 and/or [Der p 2]4 staining. Scatterplot depicting MFI ratio of allergen: streptavidin of [Der p 1]_4_-APC versus [Der p 2]_4_-APC on basophils from control and HDM-allergic samples. **(B)** Scatterplot for the proportion of CD63^+^ basophils in BAT following *in vitro* stimulation with both allergens.

## DISCUSSION

Here we present highly sensitive and specific detection of HDM sensitization by using two immunodominant HDM components as recombinant tetramers in a single CytoBas tube. The performance of Cytobas for single Group 1 (Der f 1 or Der p 1) or Group 2 (Der p 2) allergens was comparable to BAT. However, CytoBas performance was further enhanced by combined detection of both Der p 1 and Der p 2 – achievable in a single CytoBas assay – with 100% sensitivity and 96% specificity.

Several key technical differences favor the use of CytoBas over BAT. First, CytoBas does not require *in vitro* stimulation of basophils. As a result, the test is rapid, less susceptible to batch or operator differences, and can be performed on thawed, live PBMC. Second, with BAT, basophils of 5-10% of people fail to respond to *in vitro* cross-linking of IgE (non-responders), making the result uninterpretable.^45^ CytoBas has no such restrictions. Third, CytoBas is a single-tube assay and does not rely on repeat measurements for additional components or titration of the stimulus, saving time and reagents.

We here employed the baculovirus expression system in Sf21 insect cells to produce recombinant Der f 1, Der p 1 and Der p 2 proteins that were highly reactive in our BAT and CytoBas assays. Sf21 cells enable paucimannosidic glycosylation of produced proteins,^46^ which cannot be obtained in bacterial cell systems, and could be the reason for the previously observed reduced (25-fold) immunogenicity.^47^ While *P. pastoris* yeast cells are capable forming highly-glycosylated Der p 1 with allergenic activities, this was also limited in inducing an immune response.^48^ Similar to previously published plant and insect proteins by us and others,^35, 49^ we here showed that the *S. frugiperda* cells were suited for the production of recombinant HDM recombinant proteins that could be recognized by IgE of sensitized patients.

Our recombinant Der f 1 and Der p 1 proteins contained specific amino acid substitutions. Both proteins contained the C35S mutation to disrupt their protease activity, and minimize the impact on protein stability and/or cell viability in our assays.^44, 49^ To overcome the challenges of recombinant Der p 1 production, the additional S(−63)N mutation was introduced in the prodomain, disrupting a glycosylation site,^41^ which is absent in Der f 1. In line with previous reports,^35, 41^ these targeted mutations did not impair the immunoreactivity with our protein tetramers efficiently binding IgE on basophils and triggering degranulation.

CytoBas using either [Der p 1]_4_ or [Der p 2]_4_, exhibited comparable sensitivity and specificity for HDM sensitization detection as BAT. While the robustness of the ROC analysis might be affected due to the higher number of patients than controls, it does demonstrate that allergen-component binding to basophils is similarly effective for detection of allergen sensitization as *in vitro* activation. In addition, the 4 non-responders in BAT (1 patient, 3 controls) in our study were evaluable with CytoBas. Thus, basophil staining is a highly suited approach to detection functional allergen sensitization without the potential challenges of *in vitro* stimulation.

As compared to IgE serology, the CytoBas approach is more labor-intensive and costly, which can impact clinical translation. Rather than replacing standard specific IgE tests, it can be utilized for specific queries requiring differential diagnosis with 8 or more components. As these can be run in a single assay, the cost per component is drastically reduced. Furthermore, the assay can provide a same-day result within 2-3 hours in acute situations, e.g. differential evaluation of β-lactam allergy, and it provides evaluation of functional, cell bound IgE, rather than serum IgE.

All 64 patients in our study were sensitized to Der p 1 and/or Der p 2, which is consistent with previous findings that Der p 1 and Der p 2 are the highly immunodominant components of HDM (95%).^30, 50, 51^ Der p 1 and Der p 2 have been described as ‘initiator molecules’ for HDM sensitization in the context of molecular spreading,^52^ i.e. allergen sensitization arises first against these components and subsequently spreads to other components from the same allergen source.^53^ This suggests that the detection of sensitization to Der p 1 and Der p 2 can be beneficial for early screening. Furthermore, dual-sensitization to both Der p 1 and Der p 2 is associated with more concerning morbidity and a higher risk of asthma among allergic patients.^54, 55^ Based on our findings, combining Der p 1 and Der p 2 in the CytoBas approach, provides the best sensitivity and specificity for allergen detection of HDM allergic patients, reinforcing the need to use both components for future diagnoses.

Parallel evaluation of basophil staining and activation by [Der p 1]_4_ and [Der f 1]_4_ enabled investigation of the immunoreactivity between the primary sensitizer (Der p 1) and a highly homologous variant (Der f 1). All patients (n=32) who had positive staining to Der p 1 were positive for Der f 1, while 90% (n=27/30) who showed basophil activation to Der p 1 were positive for Der f 1. Der p 1 and Der f 1 amino acid sequences have a 90% similarity, and the two structures are highly homologous,^56^ so that epitopes will be largely shared between these two allergens.^57^ Still, the Der p 1 CytoBas exhibited enhanced performance capability (AUC from ROC) than Der f 1 CytoBas in our Australian patient cohort. This underpins the importance of using relevant allergen components based on geographic region^58^ to achieve reliable diagnostic results, as suggested by publications in aeroallergies^31,59^ and food allergies.^60^ The inclusion of geographically relevant allergens together, or in multiplexed form, such as combining Der f 1 and Der p 1, despite their largely shared epitopes, will increase the universality of CytoBas HDM allergy detection.

While all 64 HDM allergic patients were detected with CytoBas including Der p 1 and Der p 2, several studies have reported that 2-5% of HDM allergic patients are monosensitized to Der p 23. ^31, 61, 62^ Thus, the inclusion of other (minor) allergens, including Der p 23 and Der p 10, could be important to improve diagnosis rates. Furthermore, the inclusion of these proteins in CytoBas could inform disease prognosis. Sensitization to Der p 23 is suggested to correlate with more severe asthma symptoms and the more intense airway inflammation measured via exhaled nitric oxide (FeNO).^63^ Sensitization to Der p 10 (tropomyosin) highly relates with cross-reactivity to other tropomyosin allergens, such as shrimp and crab, which can be inducers of severe systemic anaphylaxis.^64^ The early detection of sensitization against Der p 10 can advance monitoring symptoms of ingestion of cross-reactive crustacea or snails.^65, 66^ Stratification of patients based on molecular sensitization profiles obtained from CRD^67^ can improve AIT success.^68, 69^ Therefore, it can be considered to expand the CytoBas approach beyond group 1 an group 2 HDM allergens to improve precision diagnosis and treatment decisions.^70^

Our results expand our previous findings^35^ that the CytoBas approach can be applied for multiplex component-resolved detection of aeroallergen sensitization.^36^ As the major triggers of allergic asthma and AR are HDM, grass/tree pollen, and household pets,^71–73^ a single CytoBas assay with the major components from these allergens could become an efficient option for differential diagnosis of aeroallergen sensitization.

## Author contributions

MCvZ, REO’H and MH conceived the idea for the present study. REO’H, MH and LH recruited participants and facilitated sample collection. LH, NV, MCvZ conducted experiments. PMA, BDW and PMH provided support for recombinant protein production. LH and MCvZ analyzed the data. LH and MCvZ wrote the manuscript. All authors revised and commented on manuscript drafts and approved the final version of the manuscript.

## Acknowledgments

The authors gratefully acknowledge Ms. Kirsten Deckert, Ms. Anita Hazard, Ms. Monique Dols, and Ms. Anna Mackay for the collection of clinical data and blood samples from patients; blood donation from all participants; Dr Craig Mckenzie for generous support in protocol optimization; Dr Emily Edwards for initial setup using flow cytometer; Dr Anouk von Borstel for critical reading of the manuscript. This work was supported by an NHMRC Ideas grant 2000773 and an investigator-initiated grant from Stallergenes-Greer.

## Data availability statement

The data that support the findings of this study are available from the corresponding author upon reasonable request.

