## Supplemental Table 1 for "Accurate determination of house dust mite sensitization in asthma and allergic rhinitis through cytometric detection of Der p 1 and Der p 2 binding on Basophils (CytoBas)"

**Supplemental Figures** (n=5)

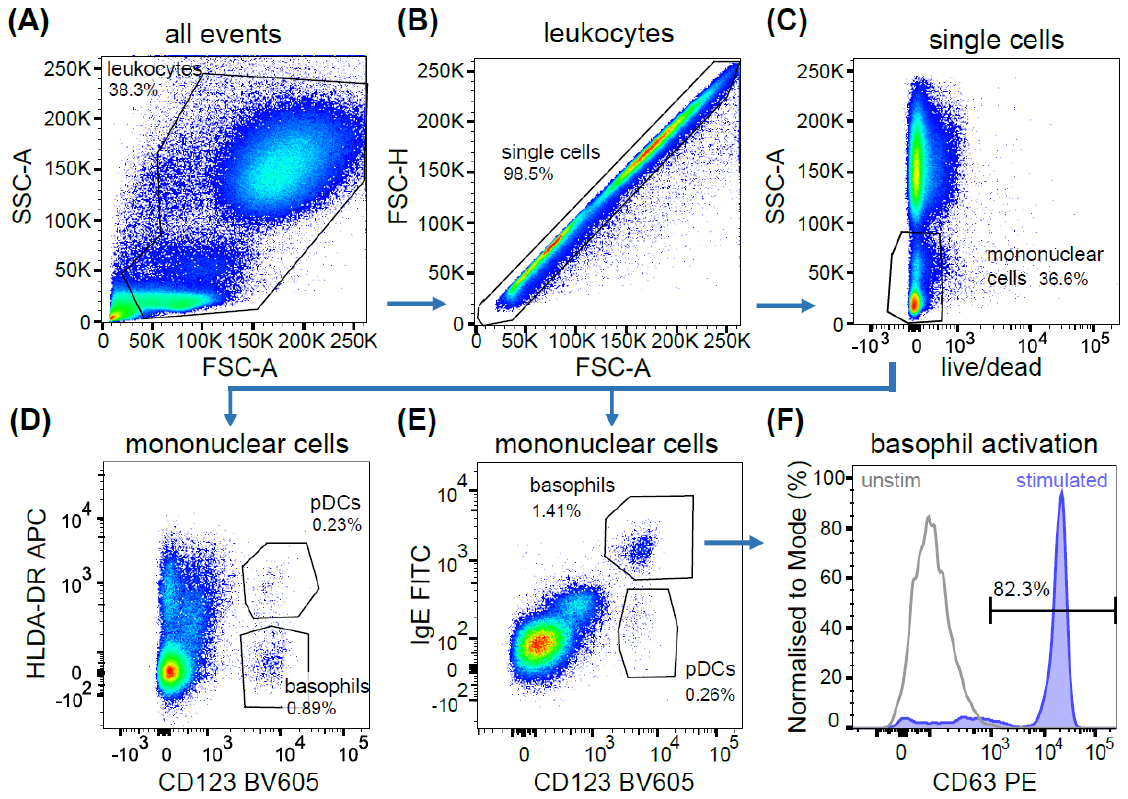
**Supp Fig 1. Gating strategy to identify activated basophils in whole blood**

Stepwise gating strategy for blood immune cells to identify (**A**) total leukocytes, (**B**) single cells and (**C**) live mononuclear cells. (**D**) Basophils are defined by the positivity for both IgE and CD123, or (**E**) by the expression of CD123 in the absence of HLA-DR for those samples in which basophils were activated with anti-IgE. (**F**) Expression of surface CD63 was used to define *in vitro* activated basophils.

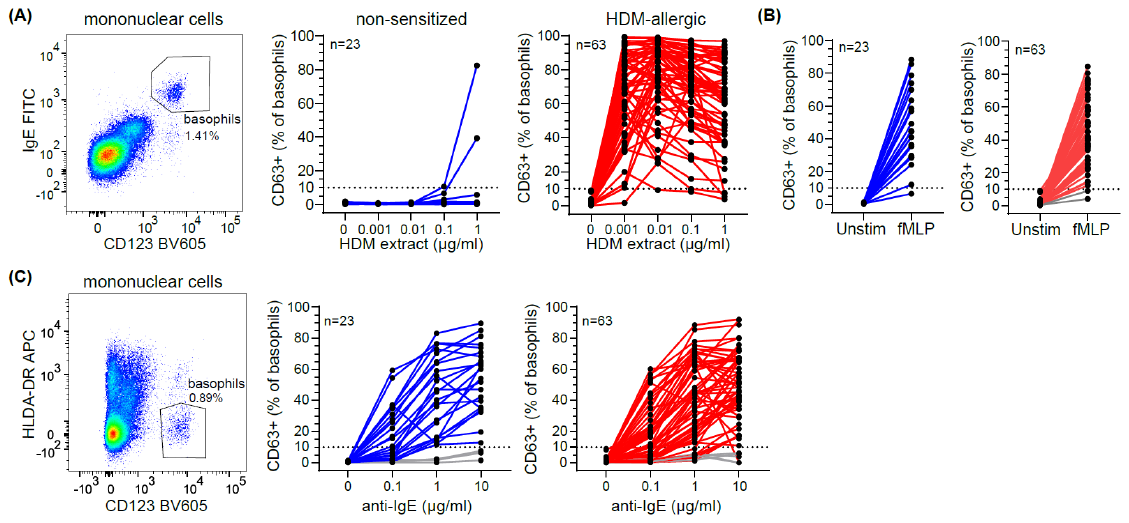
**Supp Fig 2. Basophils from both allergic and non-sensitized subjects are functionally capable of degranulation.**

**(A)** Gating strategy for the detection of basophils after stimulation with HDM extract or **(B)** fMLP in non-sensitized (blue) and HDM-allergic individuals (red). **(C)** Gating strategy using HLA-DR and CD123 for detection of basophils after anti-IgE stimulation in non-sensitized (blue) and HDM-allergic individuals (red).

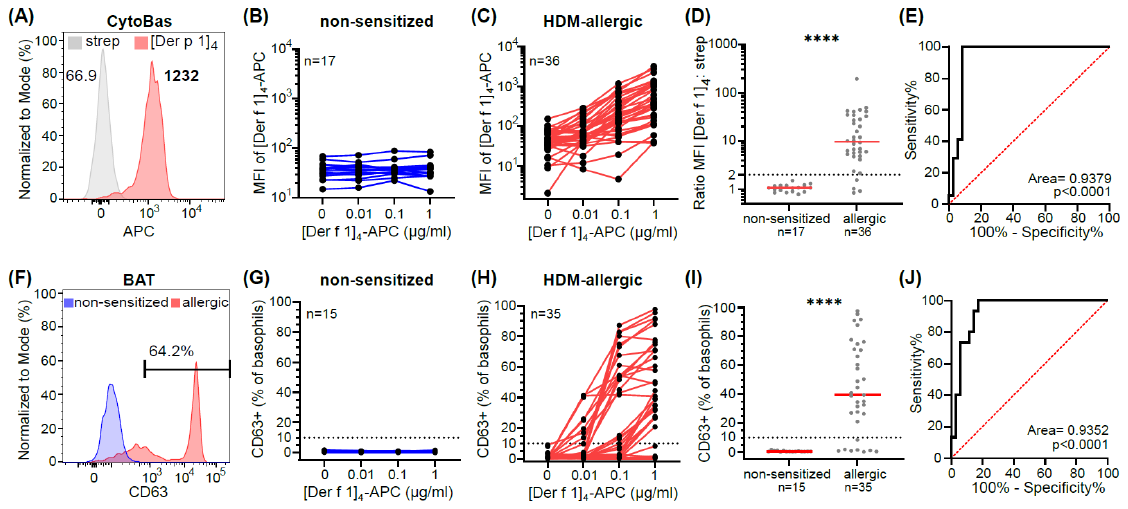
**Supp Fig 3. Performance analysis using Der f 1 tetramer in BAT and CytoBas assays.**

**(A)** Median fluorescence intensity (MFI) following stimulation of [Der f 1]_4_-APC at 1µg/mL. **(B)/(C)** frequencies of basophils from non-sensitized (blue) and allergic groups (red) following stimulation with streptavidin-APC or increasing concentrations of [Der f 1]_4_-APC. **(D)** Staining intensities of CytoBas via the MFI ratio of [Der f 1]_4_-APC at 1µg/mL over streptavidin-APC and **(E)** receiver operator curve (ROC) from MFI. **(F)** Representative CD63+ % on basophils, and **(G)/(H)** frequencies of basophils following stimulation with streptavidin-APC or increasing concentrations of [Der f 1]_4_-APC. **(I)** CD63 expression post stimulation with [Der f 1]_4_-APC at 1µg/mL and **(J)** the corresponding ROC of CD63 positivity. Statistics: Mann-Whitney U test; for ROC, Wilson/Brown method for confidence level (>95%) determination. **** p <0.0001.

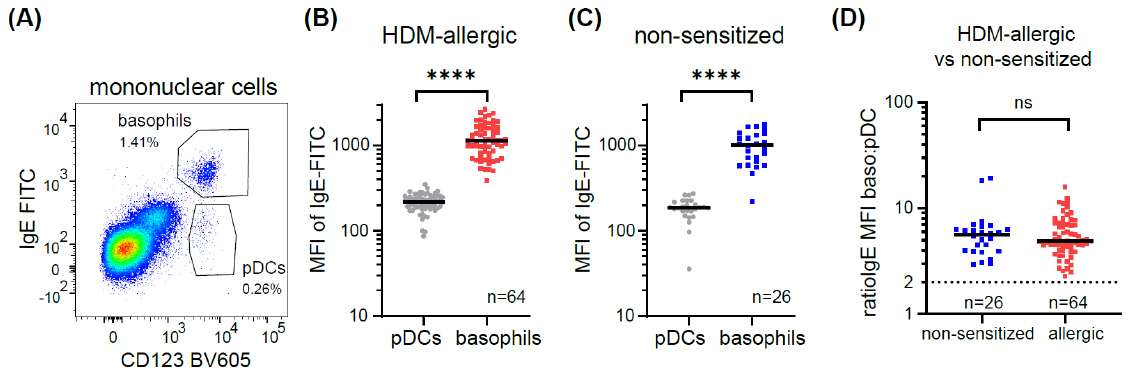
**Supp Fig 4. Surface IgE expression levels on basophils and pDCs**

**(A)** Representative flow cytometry plot of basophils and pDCs with corresponding frequencies. Median fluorescent intensity (MFI) of IgE-FITC, indicating surface IgE expression from basophils and pDC in **(B)** HDM-allergics (n=64) **and** **(C)** non-sensitized controls (n=26). **(D)** Ratio MFI of IgE staining of basophils over pDC from non-sensitized and allergic groups respectively, with median shown. Cut-off ratio of 2 is indicated by the dotted line. Statistics: Wilcoxon signed rank test. * p < 0.05, ** p < 0.01, **** p <0.0001.

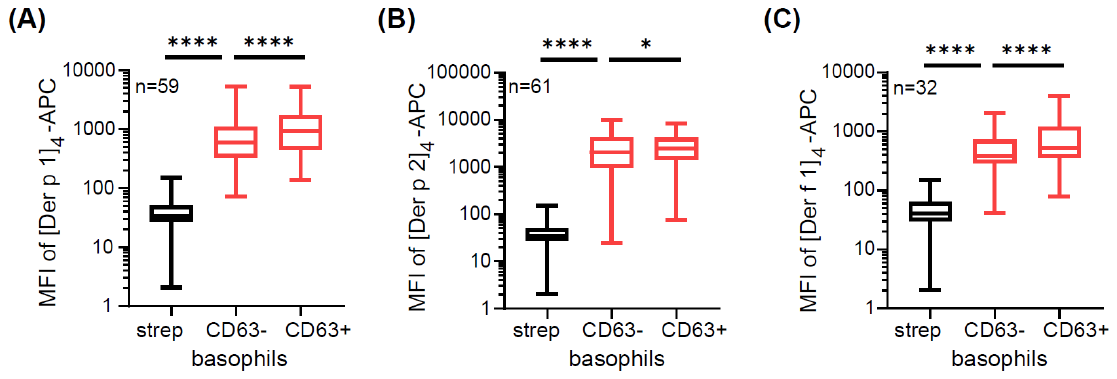
**Supp Fig 5. Allergen tetramers bind basophils in sensitized individuals irrespective of activation.**

MFI of allergen staining on CD63+ and CD63- basophils following *in vitro* stimulation with streptavidin-APC and 1 µg/ml (**A**) [Der p 1]_4_-APC, (**B**) [Der p 2]_4_-APC, (**C**) [Der f 1]_4_-APC. Statistics: Wilcoxon signed rank test. * p < 0.05, ** p < 0.01, **** p <0.0001.
